# Single cell RNA sequencing reveals mechanisms underlying a senescence-like phenotype of Alveolar Macrophages during Aging

**DOI:** 10.1101/2022.06.04.494822

**Authors:** Yue Wu, Bibo Zhu, Ruixuan Zhang, Nick P. Goplen, Xiaochen Gao, Harish Narasimhan, Ao Shi, Yin Chen, Ying Li, Haidong Dong, Thomas J. Braciale, Jie Sun

## Abstract

Alveolar Macrophages (AMs) are unique innate immune cells that reside in the alveolar space and accommodate the ever-changing needs of the lungs against internal and external challenges. During homeostasis, AMs maintain themselves through self-renewal without input from adult hematopoietic stem cells. Currently, little is known regarding the influence of aging on AM dynamics, heterogeneity and transcriptional profiles. Here, we identified CBFβ as an indispensable transcription factor that ensures AM self-renewal. Deficiency in CBFβ led to decreased proliferation and self-renewal ability of AMs. Moreover, with single cell RNA sequencing analysis of AMs from young and aged mice, we discovered that despite similarities in the transcriptome of proliferating cells, AMs from the aged mice exhibited reduced embryotic stem cell-like features. Aged AMs also showed diminished capacity for DNA repair, potentially contributing to impaired cell cycle progression and elevation of senescence markers. In accordance with the analysis, we observed reduced number of AMs in aged mice, which had defective self-renewal ability and were more sensitive to the reduction of GM-CSF. Interestingly, decreased CBFβ was observed in the cytosol of AMs from aged mice. A similar senescence-like phenotype was also found in human AMs. Taken together, we conclude that AMs in the aged host harbor a senescence-like phenotype, potentially mediated by the activity of CBFβ.

**Highlights:**

- scRNAseq revealed Alveolar Macrophage (AM) heterogeneity and self-renewal
- CBFβ is associated with AM cell cycle and facilitate AM self-renewal
- AMs displayed a senescence-like phenotype during physiological Aging
- Aging impairs CBFβ expression in mouse and human AMs

## Introduction

Alveolar Macrophages (AMs) are the major resident macrophages in the lung. AMs function as the first line of defense against microbial infections, participate in surfactant turn-over and the phagocytosis of apoptotic cells and debris while maintaining an immune suppressive state in the lung during homeostasis^1,2^. Thus, it is important to understand the mechanism underlying the maintenance of the AM compartment as well as the regulation of its function. Both human and murine AMs are derived from fetal precursors^3,4^ and upon maturation, AMs remain embryotic-stem-cell (ESC) like features of self-renewal^5^, eliminating the requirement for input from adult hematopoietic stem cells during homeostasis^6^. This notion is supported by the persistence of donor AMs in lung transplant recipients 3.5 years post-surgery^7^. To this end, AMs depend on GM-CSF and autocrine TGF-b signaling for their maturation, proliferation and maintenance^3,8,9^. Transcription factors including PPAR-g^10^, BHLHE40 and BHLHE41^11^, Bach2^12^, Hem-1^13^ as well as EGR2^14^ have been reported to be indispensable for the generation and/or maintenance of the AM compartment. Furthermore, mitochondrial-dependent oxidative metabolisms are also vital for AM self-renewal and maintenance^15,16^. However, the exact molecule program regulating AM cell cycle progression, proliferation and self-renewal remains largely undefined.

Cellular senescence describes a cellular state marked by irreversible cell cycle arrest in response to internal and external stress^17^. It is characterized by unresolved DNA damage, inability of cell cycle progression, and meanwhile, the resistance against apoptosis^18^. Cells that undergo senescence express an increased level of factors involved in cell cycle arrest, most notably p16 (encoded by *Cdkn2a*), and a secretome termed senescence associated secretory phenotype (SASP), resembling the expression of innate effector molecules at baseline^18^. Changes in cellular metabolism and the epigenetic landscape are also important components in cellular senescence^17,18^. Physiological aging lead to increased features of senescence in the lung^19^, accompanied by accumulation of pro-inflammatory cytokines, complement components and molecules indicative of oxidative stress in the alveolar lining fluid^20^. It has been shown that the senescent immune cells, particularly T cells, can independently drive tissue damage and organ aging in non-lymphoid tissues including lung tissue^19^. However, it is not known whether tissue-resident macrophages exhibit senescent-like phenotypes during aging. To this end, previous research has described a dysfunctional phenotype in AMs derived from aged host^21,22^. Nevertheless, it remains to be explored whether AMs undergo cellular senescence during aging and if so, what the exact cellular and molecular mechanisms underlying such phenomenon would be.

Core binding factor b subunit (CBFβ) is a non-DNA-binding transcription factor that binds to its DNA-binding partner, RUNX1, RUNX2 or RUNX3, to form a heterodimer which facilitates its function^23^. Core binding factors play an essential role in hematopoiesis during embryotic stage where deficiency can lead to defects in the generation of hematopoietic stem cells in the fetal liver^24,25^. Besides establishment of definitive hematopoiesis, CBFβ is also important for maturation of cells in the myeloid lineage from adult hematopoietic stem cells^26^. For instance, deficiency of a CBFβ-RUNX1 binding enhancer of IRF8 skewed the fate of myeloid progenitor cells towards differentiation into monocytes^27^. In the respiratory tract, myeloid deficiency of CBFβ caused defects in the AM compartment despite normal development of monocytes, neutrophils, DCs, and interstitial macrophages (IMs) ^28^. However, the underlying mechanisms by which CBFβ regulates AM homeostasis remains to be explored.

Here, we generated a single cell RNA sequencing (scRNAseq) dataset that comprise of a large number of AMs from young (12,358 cells) or aged mice (5,269 cells). The abundant cell number facilitated the study of AM heterogeneity with detailed resolution, enabling the identification of actively proliferating AM populations in which we explored the mechanisms governing the process of self-renewal. We observed that AMs from the aged mice exhibited reduced proliferation and self-renewal ability, displaying the features of senescence-like cells including cell cycle arrest, p16 upregulation, insensitivity to growth factor stimulation, and SASP expression. Additionally, CBFβ, a transcription factor, was found to be preferentially associated with proliferating AMs and was required for AM proliferation *in vitro* and *in vivo*. Importantly, we observed decreased CBFβ levels in AMs during aging and increased p16 expression in CBFβ-deficient AMs. Interestingly, similar senescence-like phenotype was found in AMs derived from aged humans, accompanied by a decline in expression of *CBFB* and its binding partners. Taken together, our data suggests that AMs in aged hosts undergo cellular senescence, which is partially attributed to the reduced expression of CBFβ during aging.

## Results

### scRNAseq reveals the functional heterogeneity of AMs in young host

AMs have been previously characterized with technologies that possess single cell resolution^22,29^. In order to overcome the restrained resolution of the data due to the limited number of AMs used in these studies, a large number of sorted CD11c+SiglecF+ AMs from the lungs of C57BL/6 mice at the age of 8-weeks were put through scRNAseq in this study **(Fig 1A)**. The contaminating myeloid cells, which were identified based on high expression of *Itgam* (CD11b), *Ccr2* (CCR2) and the lack of *Siglecf* (SIGLEC F), *Adgre1* (F4/80), *Mertk* (MERTK) and *Pparg* (PPAR-γ), were excluded **(Fig S1A)**. With K-nearest neighbor (KNN) graph, 6 different clusters were discovered with Louvain algorithm, a modularity optimization technique, in Seurat^30^ from the 12,385 cells that passed quality control **(Fig 1B)**. These AMs were in the same branch with trajectory analysis using Monocle3^31^, which is consistent with previous reports that AMs could maintain themselves through self-renewal during homeostasis^6^ **(Fig 1C)**. Additionally, the top 50 featured genes in each cluster **(Fig 1D)** and the enrichment scores of relevant pathways **(Fig 1E & S1B)** were utilized to obtain functional insights into each subcluster of AMs. AMs in Cluster 0 were characterized with expression of genes involved in antigen presentation and oxidative phosphorylation (OXPHOS) in high levels **(Fig 1D, S1B)**. Elevated expression of scavenger receptor genes like *Cd36* was found in cluster1, which had the highest score in the lysosome-related pathway **(Fig1 D, E)**. Such a transcriptome suggested that these AMs may be preferentially responsible for surfactant turn-over, which is further supported by the enhanced expression of genes related to lipid metabolism and antioxidants (**Fig 1D)**. Cells in cluster 2 possessed heightened expression of genes involved in intracellular or environmental stress responses **(Fig 1D)**. Cluster 3 appeared to consist of cells poised for inflammation, for they were enriched with expression of genes encoding pro-inflammatory signaling pathways, transcription factors, cytokines, and chemokines **(Fig 1D & S1B)**. AMs have been reported to possess embryotic stem cell (ESC) like features^5^, and such a profile^32^ was enriched in cluster 4 and cluster 5 - the proliferating AM clusters. Together, these data demonstrate that the AM compartment is considerably heterogeneous during homeostasis and there are different AM subclusters specialized in distinct functions including antigen presentation, surfactant turnover, inflammation, and proliferation.

**Figure 1.**
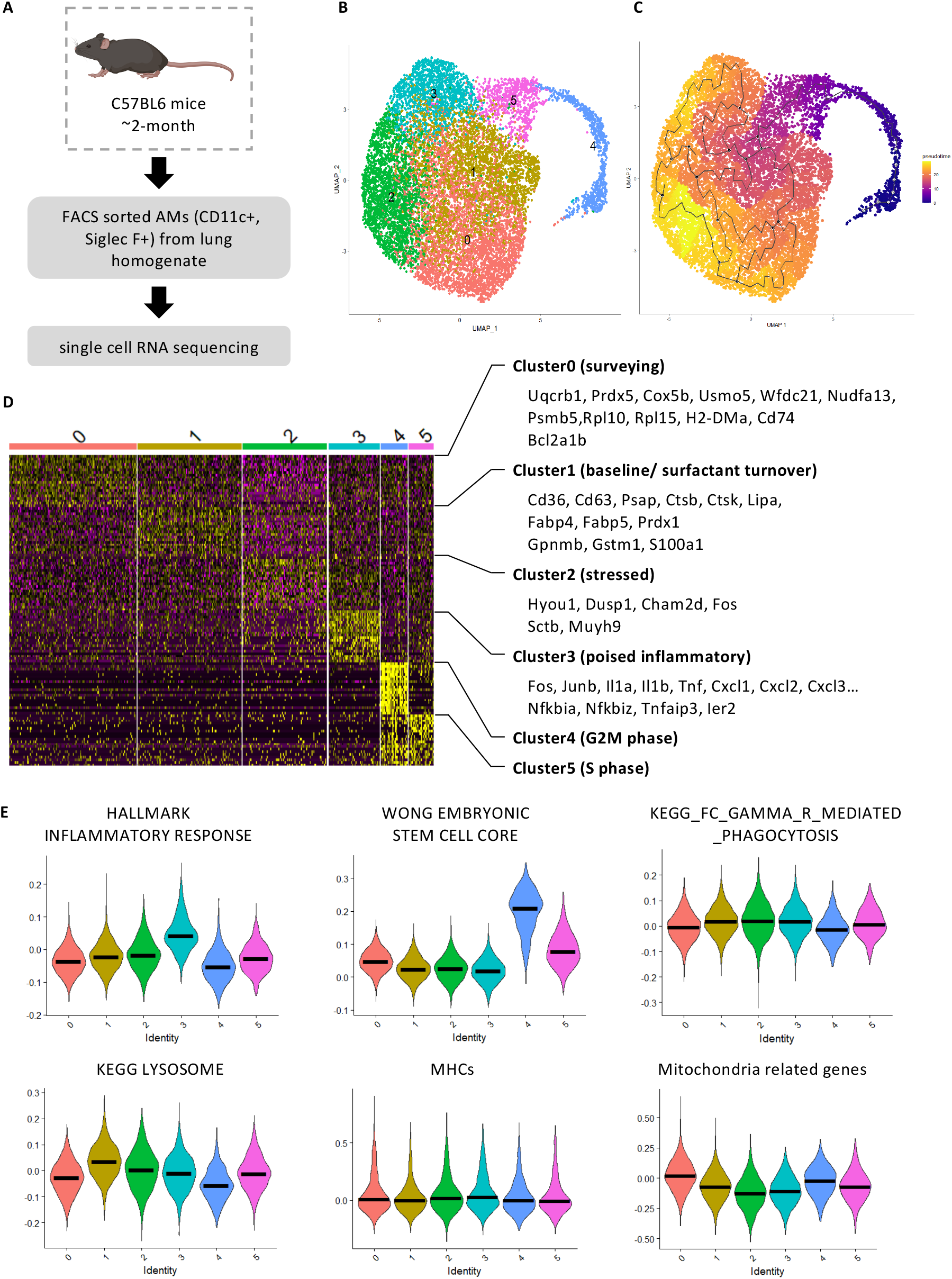
scRNAseq reveals functional heterogeneity in AMs. **A**. Schematic display the sample collection for scRNAseq. AMs (CD11c^+^ Siglec F^+^) were FACS-sorted from lung homogenate combined from multiple young mice (∼2-month-old). **B**. Visualization of the clustering of 12,358 cells with UMAP. **C**. Visualization of the trajectory analysis with UMAP. **D**. Heatmap displaying top 50 genes in each cluster with selected genes shown on the right. **E**. VlnPlot showing the module scores of gene lists of inflammatory response (MSigDB M5932), ESC-like features (MSigDB M7079), phagocytosis (MSigDB M16121), lysosome related genes (MSigDB M11266), MHC molecules and mitochondria-related genes^35^. The bar in **(E)** represent the median of the expression level of module scores of gene lists indicated among the cells in each cluster as shown in **(B)**. See also Fig S1 A and B.

### scRNAseq identifies CBFβ as a critical regulator of AM proliferation

During homeostasis, the AM compartment is maintained by self-renewal and proliferation. However, the transcriptional regulation of AM cell cycle progression remains poorly understood. To investigate the mechanisms underlying the self-renewal ability of AMs, we first identified actively proliferating cells using previously reported gene lists for G2/M or S phase markers^33^. We found that cells in cluster 4 were mainly in G2/M phase while cells in cluster 5 were mainly in S phase **(Fig 2A)**. The transcription factors enriched in cluster 4 and cluster 5 were examined to elucidate the transcriptional control of AM proliferation. Notably, the transcription factor named *Cbfb*, was found to be highly enriched in cluster 4 and 5 **(Fig.2B)**. Furthermore, *Cbfb*, but not any of the other transcription factors known to be important for AM development and/or proliferation^10–14^ corelated with the pattern of G2/M phase score and the expression of proliferation marker, *Mki67* **(Fig 2B)**.

**Figure 2.**
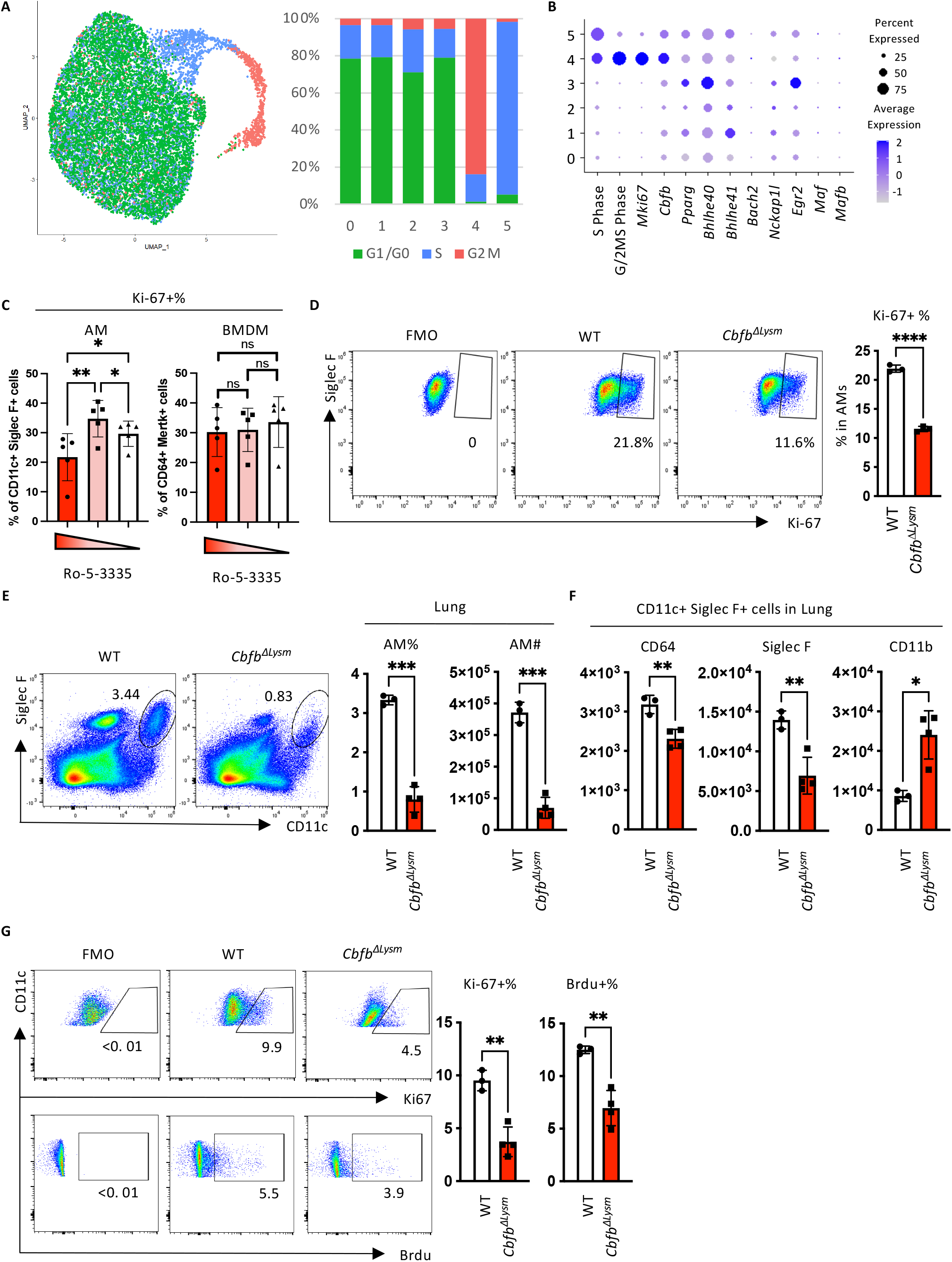
CBFβ regulates AM proliferation and self-renewal. **A**. Cell cycle analysis of AMs from the young host using list of genes associated with S and G2/M phases of cell cycle. UMAP **(left)** and quantification **(right)** of the cells predicted to be in each cell-cycle phase with respect to their identity shown in **Fig 1B. B**. Dot plot revealing the score for S-Phase or G/2M-Phase related genes, transcription factors related to proliferation and identity of AMs, and transcription factors associated with the suppression of the ESC-like feature of AMs. **C**. Percentage of Ki-67 expressing macrophages in response to its major growth factor culture upon treatment with different concentration of Ro 5-3335 (100μM, 50μM and 0 μM respectively). **D**. Adherent cells in BAL from *Cbfb*^Δ*Lysm*^ or wild-type mice were stimulated with 10ng/ml GM-CSF in vitro. The representative FACS plot **(left)** and quantification **(right)** of Ki-67 expressing AMs following 24 hours in culture. **(E to G)** Bone-marrow-chimeric mice were constructed by transferring bone marrow from *Cbfb*^Δ*Lysm*^ or wild-type (WT) mice to age and gender matched WT mice. **E**. Representative FACS plot **(left)** and quantification **(right)** of AMs in the lungs of *Cbfb*^Δ*Lysm*^ or wild-type BM chimeric mice. **F**. Geometry mean fluorescence intensity (gMFI) of markers for AM identity, CD64, Siglec F and CD11b, in AMs (defined as CD11c+SiglecF+ cells). **G**. Brdu was injected 24h before euthanization of the mice. Representative FACS plot **(left)** and the quantification **(right)** of the percentage of AMs expressing Ki67 or having Brdu incorporated in lungs from *Cbfb*^Δ*Lysm*^ or wild-type BM chimeric mice. Data were analyzed with RM one-way ANOVA with the Geisser-Grennhouse correction and multiple comparisons **(C)** or unpaired student t test with Welch’s correction **(D-G)**. Data were shown as mean ± SD. *, p < 0.05; **, p < 0.01; ***, p<0.001; ****, p<0.0001. Data in **(B)** were displayed in dots with diameter representing the percent of cells in the cluster (as shown in **Fig 1B)** that expressed the gene (or list of genes) indicated, with the depth of color indicating the average expression level. See also **Fig S1 C to E**.

CBFβ deletion led to defects in the AM compartment^28^, but the underlying mechanisms are unknown. The expression pattern of *Cbfb* suggests that it may regulate proliferation of AMs. To this end, we examined the role of CBFβ in the proliferation of AMs using *in vitro* and *in vivo* models. Since CBFβ requires RUNX family members which possess DNA-binding domains^23^ to confer its functions, we inspected data on Immgen^34^ and found that AMs mainly express *Runx1* among the RUNX family members **(Fig S1C)**. Thus, we treated AMs and bone marrow-derived macrophages (BMDMs) *in vitro* with different concentration of Ro 5-3335, a small molecule drug that inhibits the interaction between CBFβ and RUNX1, in the presence of 10ng/ml GM-CSF^16,35^. With similar viability of the cells **(Fig S1D)**, 100μM of Ro 5-3335 treatment inhibited the proliferation of AMs but not BMDMs **(Fig 2C)**. It is possible that Ro 5-3335 may have non-specific effects targeting factors other than CBFβ, we therefore examined whether genetic deletion of CBFβ resulted in diminished AM proliferation. We compared the proliferation of AMs isolated from WT or myeloid-specific CBFβ knock-out mice (*Cbfb*^Δ*Lysm*^) in response to GM-CSF stimulation *in vitro*. CBFβ-deficient AMs displayed significantly decreased proportion of Ki-67 expressing cells compared to that of WT AMs, suggesting that CBFβ is critical for AM proliferation in response to growth signals **(Fig 2D)**. We next examined whether CBFβ is important for AM proliferation *in vivo*. As *Lysz2* may also be expressed in alveolar type II cells (AT II)^36^, we ruled out potential effects of ATII CBFβ expression by generating bone-marrow chimeric mice by transferring bone marrow from *Cbfb*^Δ*Lysm*^ mice or control mice into lethally irradiated WT mice. 8 weeks post irradiation, we examined the AM compartment and proliferation following reconstitution. Deficiency in CBFβ led to decreased number of AMs in the lung **(Fig 2E)** and BAL **(Fig S1E)**. The AMs also exhibited decreased expression of macrophage marker, CD64, and immature AM phenotype, evidenced by decreased expression of Siglec F and increased expression of CD11b **(Fig 2F)**. Consistent with our *in vitro* findings, the proliferation, as shown by Ki-67 expression or Brdu incorporation, was decreased in AMs lacking CBFβ **(Fig 2G)**. Taken together, we identify CBFβ as a potential important transcription factor for AMs’ identity and self-renewal ability.

### scRNAseq distinguish the phenotype of AMs from young and aged mice

To explore how aging influences the phenotype of AMs and the underlying molecular mechanism, scRNAseq data of AMs from young (∼2-month-old) and aged (∼22-month-old) mice were combined for analysis **(Fig 3A)**. The same number of cells were put into analysis in order to ensure equal input from both groups in the detection of differential expressed genes. Moreover, other myeloid cells (cluster 10) which express high level of *Itgam* (CD11b) and *Ccr2* (CCR2) as well as low level of *Siglecf* (Siglec F), *Adgre1* (F4/80), *Mertk* (MERTK) and *Pparg* (PPAR-γ) were removed **(Fig S2A)**. Using the same method described previously, re-clustering of the remaining cells as well as further cell cycle and trajectory analysis were then performed **(Fig 3C&D)**. Cluster 6 was enriched in a gene profile of cells in S phase, while cluster 8 and 10 were enriched in a gene profile of cells in G2/M phase **(Fig 3B & C)**. Because of the self-renewing nature of AMs, the proliferating cells were chosen as “stem cells” for trajectory analysis. Although “upstream” proliferating cells (clusters 6, 8, 10) had similar transcriptional profile, AMs from young and aged mice gradually branched into 2 different streams with cluster 0 or cluster 4 at the end of the AMs from aged mice or young mice, respectively. **(Fig 3B&D)**. Both cluster 0 and cluster 4 can be defined as poised inflammatory cells since both proinflammatory and anti-inflammatory genes were enriched **(Fig S2B&D)**.

**Figure 3.**
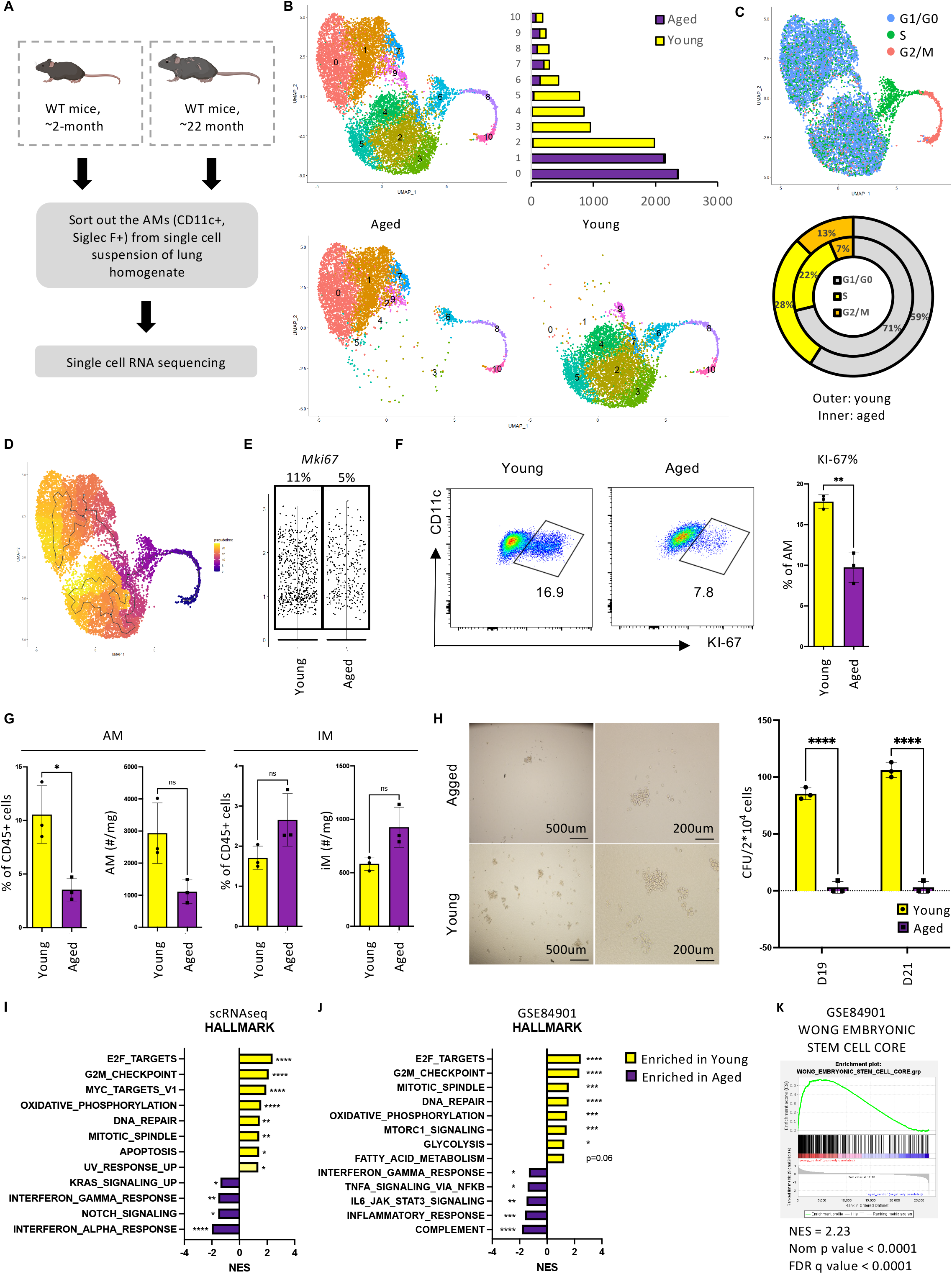
AMs from the aged mice display reduced self-renewal ability. **A**. Schematic display the sample collection for scRNAseq. AMs (CD11c^+^ Siglec F^+^) were FACS-sorted from lung homogenate combined from multiple young (∼2-month-old) or aged (∼24-month-old). **B**. Visualization of the clustering of 10,531 cells with UMAP displaying over-all clustering **(upper left)** and the quantification of cell number in each cluster **(upper right)** as well as split view with regard to their hosts **(lower). C**. Cells were denoted to indicated cell cycle phase based on PCA analysis using genes sets feature of S phase and G2M phase, as displayed in UMAP **(top)** and quantification of cells in each phase **(bottom). D**. UMAP showing the trajectory analysis using Monocle3 with the proliferating cells as “stem cells”. **E**. Violin plot displaying *Mki67* expression in AMs of young or aged mice from scRNAseq. Quantification of percentage of cells expressing *Mki67* was shown on the top. (**F and G)** Flow cytometry of macrophages in the lung of young and aged mice. **F**. Percentage of AMs expressing Ki-67 from the lung of young and aged mice shown by representative FACS plot **(left)** and quantification **(right). G**. Quantification of percentage and absolute number of AMs **(left)** and IMs **(right). H**. Colony formation assay using AMs from young and aged mice. Demonstrated by representative pictures of the colony **(left)** and quantification of colony unit number **(right). I**. Bar graph showing selected results of pathway analysis comparing AMs from young and aged mice using GSEA. **J**. Bar graph showing selected results of pathway analysis comparing AMs from young and aged mice in published microarray data (GSE84901 using GSEA). **K**. GSEA comparing the ESC like features (MSigDB M7079) of AMs from young and aged mice in published microarray data (GSE84901). Data were analyzed with unpaired student t test with Welch’s correction **(F, G and H)**. Data were shown as mean ± SD. *, p < 0.05; **, p < 0.01; ***, p<0.001; ****, p<0.0001. See also **Fig S2**.

Clusters 2, 3, and 4 mainly derived from young mice, and followed an order of 3 => 2 => 4 in pseudotime **(Fig 3D)**. Cluster 3 had a higher score in ESC-like features **(Fig S2D)** and expressed transcription factors known to facilitate anti-apoptotic functions, i.e., Bcl2a1b and Bcl2a1d **(Fig S2B)**. Moreover, cluster 3 was enriched in mitochondria related profile and OXPHOS related genes **(Fig S2C)**, which is consistent with previous reports that OXPHOS is required for the proliferative function of AMs^16,35^. Cluster 2 appeared to be an intermediate stage between cluster 3 and cluster 4. Meanwhile, clusters representing AMs from the aged mice, cluster 0 and cluster 1, were enriched in genes for MHC II molecules **(Fig S2D, E)**. Together with the trajectory analysis **(Fig 3D)**, cluster 1 likely represents AMs differentiating from cluster 0. In relatively upstream of the pseudotime **(Fig 3D)**, cluster 7 and cluster 9 were enriched in genes associated with lysosome activity and lipid metabolism **(Fig S2B&C)**, suggesting their preferential role in surfactant turnover. Lastly, AMs in cluster 5 may undergo apoptosis as they were enriched solely in genes for cytoskeleton compartments and genes indicative of mitochondria leakage like mt-Atp8 **(Fig S2B)**. Taken together, the combined dataset of AMs from young and aged mice revealed functional heterogeneity in AMs, with marked distinction in single cell transcriptional profiles with age.

Consistent with previous reports that AMs in the aged hosts exhibited decreased ability for self-renewal^21,22^, the proportion of cells denoted to the S phase or G2/M phase decreased in the aged hosts **(Fig 3C)**. This is confirmed by the decreased percentage of cells expressing *Mki67* during aging in the combined scRNAseq data **(Fig 3E)**, which was further validated by flow cytometry **(Fig 3F)**. The reduced proliferation of AMs led to a significant reduction in number of AMs but not IMs in the lungs of aged mice **(Fig 3G)**. To directly evaluate the self-renewal ability of AMs in addition to proliferation, we performed the colony-formation assay, where AMs from young and aged mice were sparsely seeded on matrix gels and the colonies derived from these cells were examined *after in vitro* culture. AMs from aged hosts displayed decreased number and size of colonies **(Fig 3H)**, suggesting diminished potential for self-renewal. Consistently, pathway analysis using the gene set enrichment analysis (GSEA)^37,38^ of the transcriptome of 1000 cells randomized out from the scRNAseq and published microarray data of AMs from young and aged hosts (GSE84901)^21^ confirmed that AMs from young hosts were enriched in the proliferation-related transcriptome including pathways related to cell cycle checkpoints, cell replication machinery, DNA repair, and telomere capping **(Fig 3I, J & S3B)**. Furthermore, we found that AMs from aged mice exhibited diminished expression of genes involved in ESC-like features **(Fig 3K & S3A)** ^32^. Taken together, these data demonstrate that aging leads to a reduction of ESC-like features and self-renewal ability of AMs.

### AMs in the aged mice exhibit senescence-like phenotype

Cellular senescence describes a state of permanent cell-cycle arrest together with increased senescent markers and senescence-associated secretory phenotype (SASP), characterized by baseline secretion of cytokines and chemokines^17,18^. Aside from the decreased self-renewal ability, AMs in aged hosts have decreased apoptosis related profile and enhanced inflammation related profile **(Fig 3I, J & Fig S3B)**. We thus hypothesize that AMs from aged hosts possess a senescence-like phenotype. Indeed, although we were not able to observe a specific subset of AMs from the aged mice that possesses the senescence-like phenotype, the bulk AMs from aged hosts showed elevated expression of the reported list of genes associated with senescence^39^ **(Fig 4A)**. To further confirm such phenotype, markers associated with cellular senescence, which included genes that emphasize cell-cycle regulation and inflammation, were analyzed in the scRNAseq data. With regards to cell-cycle regulation related markers, aging led to increased *Cdkn2a* (p16) but not *Cdkn2d* (p19), *Cdkn1a* (p21), or *Trp53* (p53) expression in AMs **(Fig 4B)**, which was validated by qPCR **(Fig S3C)**. Moreover, AMs from aged mice displayed decreased proliferation, evaluated by the proportion of cells expressing Ki-67, upon in vitro stimulation with GM-CSF **(Fig 4C and D)**. Even upon saturation (10ng/ml) of GM-CSF, AMs from aged mice had fewer Ki-67 expressing cells, indicating that a significant proportion experienced cell cycle arrest **(Fig 4C)**. When normalized to the 10ng/ml group in AMs from young and aged mice, AMs from aged mice were more sensitive to the drop of GM-CSF concentration **(Fig 4D)**. Increased expression of other senescence-related markers including *Serpine1* (PAI1) and *Mmp8* (MMP8) was also observed **(Fig 4B)**. Both PAI-1 and MMP8 have been implicated in the pathogenesis of lung fibrosis^40,41^, and TGF-β1/p53/PAI-1 signaling cascade has been reported to be involved in cellular senescence^42–44^. Enhanced lysosomal activity, indicated by elevated senescence-associated β-galactosidase (SA-β-gal) expression, has been considered to be a hallmark for cellular senescence^45,46^. SPiDER-βGal (a dye that enables staining of live cells) staining confirmed that AMs in the aged lung were undergoing cellular senescence **(Fig 4E**). It has been reported that the senescent cells promote the development of lung fibrosis^47^, and the accelerated immunosenescence mouse model has suggested a potential role of immunosenescence in solid organ aging^19^. Previous studies have shown collagen accumulation in the lungs of both mice^48^ and humans^49^. As the major immune cell population in the lung during homeostasis, AMs from aged hosts have increased expression of mRNA for Collagen IV and Collagen XIV **(Fig S3E)**, raising the possibility of direct collagen deposition in the aged lung^50^.

**Figure 4.**
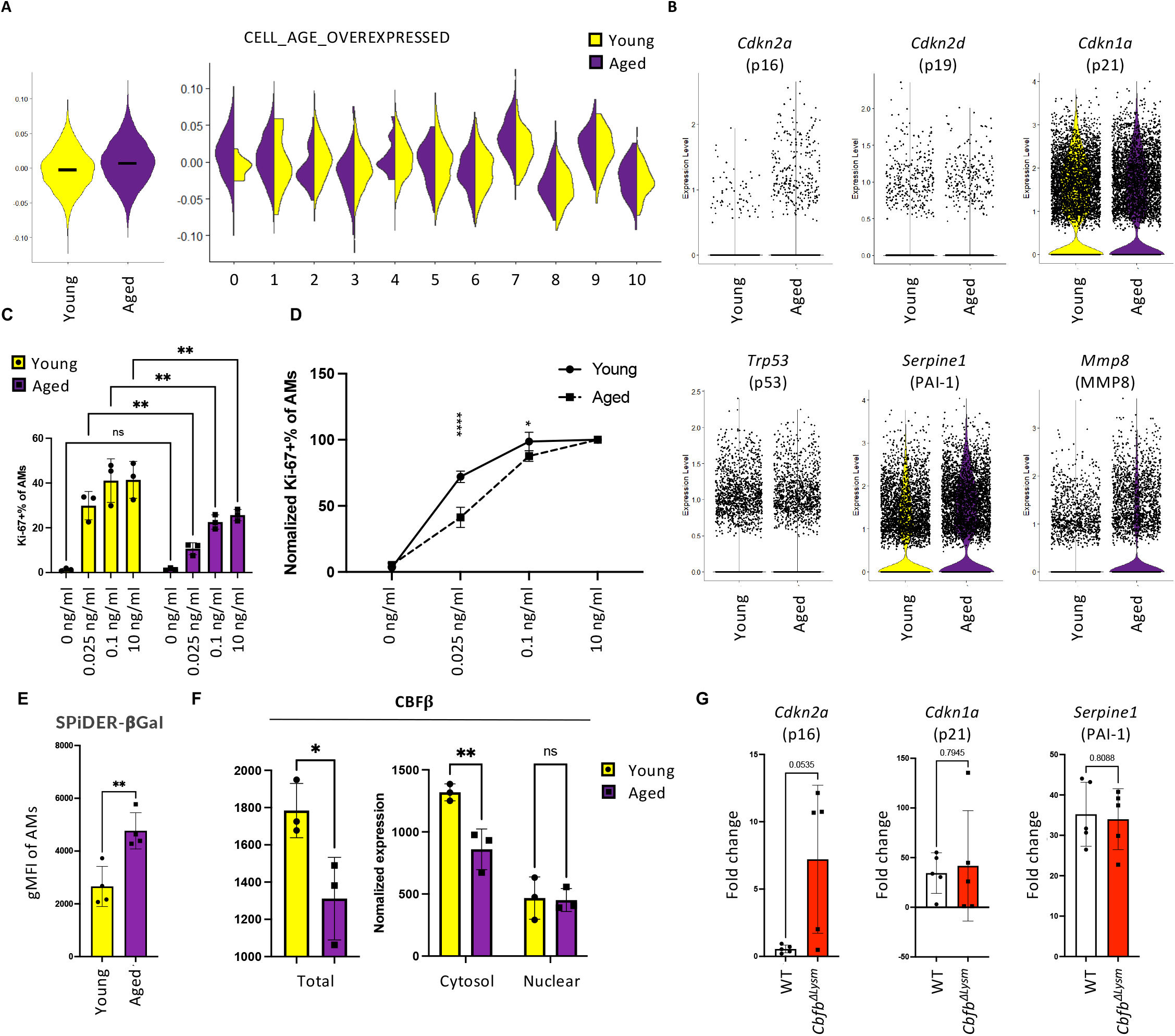
AMs from the aged mice possess a senescence-like phenotype. **A**. Module score of the list for genes that has been shown to be associated with cellular senescence was calculated for the AMs^39^. Violin Plot comparing the score between AMs from young and aged mice **(left)**, and the score among the clusters as shown in **Fig 3B (right). B**. Violin plots displaying selected senescence markers with gene names and protein names shown on the top respectively. (**C and D)** AMs were isolated from young and aged mice, followed by in vitro culture with different concentration of GM-CSF. **C**. Bar graph displaying the percentage of Ki-67 expressing AMs upon *in vitro* stimulation with indicated concentration of GM-CSF. **D**. The percentage of proliferating cells were normalized with that upon saturated concentration of GM-CSF (10ng/ml) of AMs from young or aged mice. **E**. Geometric mean of Fluorescence Intensity(gMFI) of SPiDER-βGal staining in AMs from lung homogenate of young or aged mice. **F**. Quantification of CBFβ in the cells in the BAL from young and aged mice with Western Blot. The protein level was normalized to the total protein stain. **G**. qPCR was performed with the cells in the BAL from *Cbfb*^Δ*Lysm*^ or control mice. Relative expression of selected markers for cellular senescence was displayed. Data were analyzed with Two-Way ANOVA with multiple comparisons **(C, D and F, right)** and unpaired student t test with Welch’s correction **(E, F, left and G)**. Data were shown as mean ± SD. *, p < 0.05; **, p < 0.01; ****, p<0.0001. The bar in **(A)** represent the median of the expression level of module scores of the gene list of cellular-senescence-associated genes among the cells comparing AMs from young and aged hosts^39^. See also **Fig S4**.

To further investigate the potential molecular mechanisms underlying this phenotype, the scRNAseq of AMs from young and aged mice was explored. The expression of growth factor receptors, autocrine *Tgfb1*^9^, and transcription factors known to be important for the proliferation of AMs were similar in AMs from young and aged hosts **(Fig S4A and B)**. Moreover, the expression levels of *Maf* (cMAF) and *Mafb* (MAFB), two transcription factors considered to be inhibitory to the ESC like features of AMs, were found to be similar between the groups^5^ **(Fig S4C)**. Given the potential role of CBFb in promoting the proliferation of AMs from young mice, the role of CBFb in the senescence-like phenotype observed in AMs from the aged hosts was evaluated. There were no significant differences in the expression of *Cbfb* and its binding partners between AMs from young and aged mice **(Fig S4E and F)**. However, the protein level of CBFb in AMs from the aged mice was significantly lower than that of the young mice **(Fig4F and Fig S4G)**, suggesting that reduced CBFb protein expression may contribute to the diminished proliferation and increased senescence-related features observed in AMs from aged mice. In support of this idea, deficiency of CBFb in AMs led to an increase in *Cdkn2a* (p16), but not *Cdkn1a* (p21) or *Serpine1* (PAI1) **(Fig 4G)**. Taken together, these data suggest that impaired CBFb expression may at least partially contribute to diminished proliferation and increased senescence in AMs during aging.

### scRNAseq revealed conserved phenotype in AMs from human

So far, we have described the senescence-like phenotype in AMs during aging in mice. In order to investigate whether a similar phenotype would be observed in AMs from humans, published scRNAseq on AMs from rejected organ donors (GSE 128033) was extracted and re-analyzed^51^ **(Fig S5A and B)**. Individuals were separated into young (<40 years old), middle-aged (40-60 years old) and aged (>60 years old) groups. In accordance with the observations in mice, aging led to a decrease in the proportion of *MKI67* expressing AMs **(Fig 5A)**. With increasing age, AMs in human lungs showed decreased ESC-like features as well as increased senescence-related and inflammatory profiles **(Fig 5B)**. Importantly, we observed the enrichment of *CBFB* in proliferating populations **(Fig 5C)**, consistent with *Cbfb* expression patterns in mouse AMs. Of note, we observed downregulation of *CBFB* and *RUNX1* mRNA expression during aging in human AMs **(Fig 5D)**. Moreover, the mRNA levels of certain growth factor receptors (*CSF2RB, TGFBR1, TGFBR2*), autocrine *TGFB1*, and proliferation related transcription factors (*BHLHE41, NCKAP1L* and *EGR2*) decreased in aged individuals **(Fig S5C and D)**. Elevation of *MAFB* was also observed **(Fig S5E)**, in addition to increased expression of *CDKN1A* (p21) but not *CDKN2A* (p16) in AMs from aged individuals **(Fig 5E)**. Thus, AMs exhibited diminished proliferation and increased cellular senescence during aging in humans as well, although underlying mechanisms may differ from those of mice.

**Figure 5.**
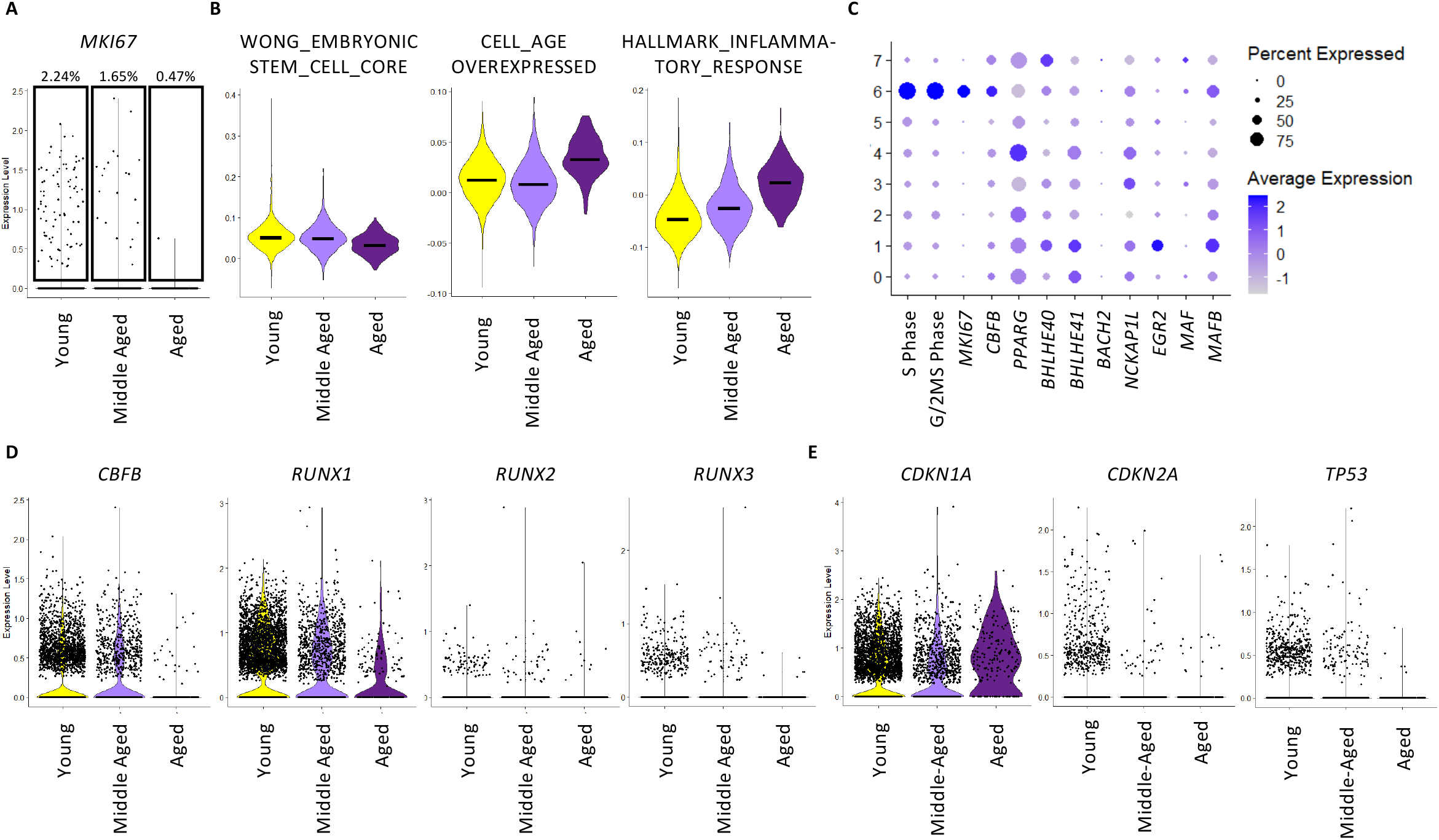
AMs from the elderly possess a senescence-like phenotype. Data of human AMs were extracted from the scRNAseq of lung donor candidates (GSE 128033). Individuals involved were divided into young (<40 years old), middle-aged (40-60 years old) and aged (>60 years old) group. **A**. Violin Plot displaying the *MKI67* expression in each group. Quantification of *MKI67* expressing cells was shown on the top. **B**. Violin Plots demonstrating the module scores of lists of genes that associated with ESC-like features (MSigDB M7079), cellular senescence^39^ and inflammatory response (MSigDB M5932). **C**. DotPlot revealing the score for S-Phase or G/2M-Phase related genes, transcription factors related to proliferation and identity of AMs, and transcription factors associated with the suppression of the ESC-like feature of AMs. **D**. Violin Plots displaying mRNA level of *CBFB* and RUNX family members (*RUNX1, RUNX2*, and *RUNX3*). **E**. Violin plots revealing selected markers for cellular senescence. The bar in **(B)** represent the median of the expression level of module scores of gene lists indicated among the cells in each cluster as shown in **(Fig S5B)**. Data in **(C)** were displayed in dots with diameter representing the percent of cells in the cluster (as shown in **Fig S5B)** that expressed the gene (or list of genes) indicated, with the depth of color indicating the average expression level. See also **Fig S5**.

## Discussion

In this study, the scRNAseq library with abundant cell number ensures the statistical strength needed to reveal functional heterogeneity within AMs. The transcription factor, CBFβ, was identified to be associated with AMs undergoing active cell cycling and was essential for AM proliferation and self-renewal. We then described the senescence-like phenotype in AMs from aged mice and humans. Moreover, our data revealed that impaired CBFβ expression during aging at least partially contributes to the observed phenotype.

Aging has been associated with decreased proliferation of AMs, resulting in reduced AM numbers^21,22^. Consistent with these studies, we observed decreased ESC-like features as well as abrogated proliferation and self-renewal of AMs during aging. However, the mechanisms underlying the acquisition of this phenotype during aging remains unclear. *McQuattie-Pimentel et al*. has attributed this to the aging lung environment^22^. Indeed, it has been shown that Alveolar type II (AT II) cells could promote the proliferation of AM by providing GM-CSF^52^ and through synchronization of Ca^2+^ flow^53^. The lung environment could also regulate AM function, namely the cell cycle regulatory genes like *Ccnd1* and *Mdm2*, through epigenetic alterations^54,55^. Moreover, the pro-inflammatory molecules in the alveolar lining fluid^20^ may promote the pro-inflammatory phenotype of AM even during homeostasis in physiological aging. However, in the bulk-RNA-sequencing data of AMs that had been adoptively transferred from aged mice into young mice or vice versa, it was revealed by k-means clustering of the differential expressed genes that one cluster of genes remains unchanged despite the change of lung environment^22^. Such phenotype suggested that there may exist a cell autonomous mechanism involved in the defect of AM proliferation from the aged hosts. Additionally, one study utilizing a *Ms4a3-*Cre-driven lineage tracing model has reported the accumulation of monocyte derived macrophages (MoM) in the pool of AMs during homeostasis^56^. In the same report, it was shown that MoM could composing up to ∼60% of total AMs at the age of 36-week-old. However, such data were not supported by long-term parabiosis (5 months of pairing)^6^ and bone-marrow-chimera with thoracic shield experiments (∼20 months post reconstitution)^22^. Constant challenges, i.e. long-term exposure to PM 2.5, experienced by the human lung throughout life could “open” the “niche” for AMs. Although there is a lack of studies on the “dirty” mouse models during aging for mimicry, it would be conceivable to consider the differences potentially brought by ontology in such setting. Taken together, it is possible that both cell autonomous and non-autonomous factors could contribute to the declined self-renewal ability of AMs during aging, but the exact mechanism remains to be explored.

Enhanced cellular senescence has been described in the lungs of both aged mice^57^ and humans^49^. Previous reports attributed the accumulation of senescent cells in the lung predominately to the stromal cell compartment^48^. Our data suggest that AMs, the self-renewing resident macrophage, also undergo cellular senescence during aging. Specifically, we have shown that AMs from aged mice displayed cell cycle arrest, decreased profile for apoptosis, enhanced pro-inflammatory profile at baseline, and increased expression of senescence markers, i.e., the SA-β-gal staining. The concept of AMs being senescent is controversial since they could be activated in the pro-inflammatory alveolar lining fluid^20^. However, besides the potential cell autonomous mechanism driving the abrogated proliferation discussed above, our *in vitro* experiments indicated that even under the same environment, AMs from aged mice were less sensitive to GM-CSF stimulation. Furthermore, they displayed decreased self-renew ability in the colony formation assay, confirming a potential senescence-like phenotype. Notably, previous studies has suggested that senescent fibroblasts^47,48^ and ATII cells^42^ may contribute to increased collagen deposition in aged lungs. However, senescence in immune cells may also be important for the aging of lung^19^. For instance, adoptive transfer of the splenocytes from aged, but not young mice could induce enhanced senescence in the lung of recipient mice^19^, suggesting a prominent role for aged immune cells in shaping the aged lung environment. However, the presence of senescent immune cells and their physiological role in the lung remains to be explored. Since AMs represent the primary immune cells present in the lung during homeostasis, it is plausible to assume that the senescence of AMs may contribute to increased lung tissue fibrosis during aging. Such a possibility warrants further studies. Furthermore, we found that AMs from aged mice exhibited increased expression of *Col4a1, Col4a2*, and *Col14a1* when compared to young mice. Thus, aged AMs may also directly contribute to lung fibrosis through the deposition of collagen.

Understanding the mechanisms underlying the dysfunctional phenotype of AMs during aging may help to identify factors and therapeutic targets to promote healthy aging of the lung. Our data suggests that impaired CBFβ expression in AMs may at least in part, contribute to the diminished proliferation and enhanced cellular senescence in AMs during aging. Thus, augmenting CBFβ expression and/or its transcriptional activities may facilitate AM self-renewal and mitigate the development of senescence during aging. Such future studies would require clear understanding of the factors that induce CBFβ expression in AMs and mechanistic basis of impaired CBFβ expression during aging.

## Limitations of the study

Although we described the senescence-like phenotype of AMs during aging, we were not able to address the acquisition of this phenotype. Furthermore, our studies also did not address the physiological function of AM senescence in the lung during aging. Additionally, since the sample size of the human scRNAseq is limited, whether human AMs definitively undergo the same cellular senescence phenomena requires future validation.

## Figure Legend

**Figure S1 Characterization of the transcriptome and the role of Cbfβ in AMs**.

**A**. Violin Plots displaying markers to distinguish AMs from other contaminating myeloid cells in the scRNAseq data before subsetting for AMs. **B**. Visualization of the module scores of genes lists concern cellular metabolism with respect to the clusters shown in **Fig1 B.C**. Expression level of *Cbfb, Runx1, Runx2*, and *Runx3* using data from Immgen. **D**. Percentage of live cells of AMs or BMDMs upon in vitro treatment of different dose of Ro 5-3335 (100μM, 50μM and 0 μM respectively). **E**. Quantification of percentage and absolute number of AMs in the BAL in bone marrow chimera mice with bone morrow from *Cbfb*^Δ*Lysm*^ or WT mice.

Data were analyzed with unpaired student t test with Welch’s correction **(E)**. Data were shown as mean ± SD. **, p < 0.01; ****, p<0.0001. The bar in **(B)** represent the median of the expression level of module scores of gene lists indicated among the cells in each cluster as shown in **(Fig 1B)**.

**Figure S2 scRNAseq reveals heterogeneity of AMs from young and aged mice**.

**A**. Violin Plots displaying markers to distinguish AMs from other contaminating myeloid cells in the scRNAseq data before subsetting for AMs. **B**. Heatmap showing top50 genes featured in each cluster (shown in **Fig 3B**) with selected genes shown on the right. **C**. Dot plot demonstrating the module score using selected gene lists. **D**. Violin plots exhibiting module score of selected gene lists of inflammatory response (MSigDB M5932), phagocytosis (MSigDB M16121), MHC molecules and ESC-like features (MSigDB M7079) in each cluster. **E**. Average expression level of MHC molecules in AMs from young and aged mice.

Data were analyzed with unpaired student t test with Welch’s correction **(E)**. Data were shown as mean ± SD. ****, p<0.0001. The bar in **(C)** represent the median of the expression level of module scores of gene lists indicated among the cells in each cluster as shown in **(Fig 3B)**.

**Figure S3 Pathway analysis revealed senescence-like phenotype in AMs from aged mice**.

**A**. Violin Plot displaying module score for ESC-like features (MSigDB M7079) in AMs from young and aged mice. **B**. Bar graph showing selected pathway analysis results using gene lists from Gene Ontology comparing AMs from young and aged mice with microarray data (GSE840901). **C**. Quantification of relative expression of selected markers for cellular senescence evaluated by qPCR. **D**. Violin plots comparing the mRNA expression for Collagen IV and XIV in AMs from young and aged mice.

Data were analyzed with GSEA **(B)** and unpaired student t test with Welch’s correction **(C)**. Data were shown as mean ± SD. *, p<0.05; **, p < 0.01; ***, p<0.001; ****, p<0.0001. The bar in **(A)** represent the median of the expression level of module scores of the gene list for ESC-like features (MSigDB M7079) comparing AMs from young and aged hosts.

**Figure S4 Characterization of proliferation related profile in AMs from young and aged mice**.

**(A-E)** Data are from scRNAseq of AMs from young and aged mice as shown in **Fig3. A**. Violin plots showing expression level of mRNA for GM-CSF receptors, TGF-β receptors, M-CSF receptor, and autocrine TGF-β. **B**. Violin plots displaying transcription factors associated with the proliferation and identity of AMs. **C**. Violin plots demonstrating mRNA level of transcription factors that suppress the ESC-like features of AMs. **D**. Dot plot revealing the score for S-Phase or G/2M-Phase related genes, transcription factors related to proliferation and identity of AMs, and transcription factors associated with the suppression of the ESC-like feature of AMs. **E**. Violin plots displaying mRNA level of *Cbfb* and RUNX family members (*Runx1, Runx2*, and *Runx3*). **F**. Quantification of relative mRNA level of *Cbfb* in AMs from young and aged mice using qPCR. **G**. Western Blot of the bands for CBFβ **(top)** and total protein stain **(bottom)**.

Data were analyzed unpaired student t test with Welch’s correction (F). Data in **(D)** were displayed in dots with diameter representing the percent of cells in the cluster (as shown in **Fig 3B)** that expressed the gene (or list of genes) indicated, with the depth of color indicating the average expression level.

**Figure S5 Characterization of proliferation related profile in AMs from human**.

Data of human AMs were extracted from the scRNAseq of lung donor candidates (GSE 128033). Individuals involved were divided into young (<40 years old), middle-aged (40-60 years old) and aged (>60 years old) group. **A**. Dot plot showing markers for validation of the identity of AMs. **B**. Identification of the clusters using the integrated data of all samples, as shown in UMAP **(left)**. Same data was demonstrated with respect to age group **(middle)** and original sample id in GSE 128033 **(right). C**. Violin plots showing expression level of mRNA for GM-CSF receptors, TGF-β receptors, M-CSF receptor, and autocrine TGF-β. **D**. Violin plots displaying transcription factors associated with the proliferation and identity of AMs. **E**. Violin plots demonstrating mRNA level of transcription factors that suppress the ESC-like features of AMs.

Data in **(A)** were displayed in dots with diameter representing the percent of cells in the cluster **(B)** that expressed the gene indicated, with the depth of color indicating the average expression level.

## Materials and Methods

### Method details

#### Mice and bone marrow chimera

Female aged C57BL/6 mice were received from National Institutes of Aging at the age around 20-21 month. Their young control, around 2-month-old, were bred in house from breeders set up with C57BL6 mice purchased from Jackson Laboratory (Harbor, ME). Beddings of the cages harboring young and aged mice were exchanged once per week for at least 4 consecutive weeks to ensure the homogenization of microbiota. The *Cbfb*^Δ*Lysm*^ mice were obtained from Dr. Thomas J. Braciale (UVA).

To generate bone marrow chimera, bone marrow cells were isolated from Cbfb^ΔLysm^ or Control mice as described previously ^58^. In brief, after long bones were isolated and cleaned, the ends of these long bones were cut open and put facing down into a 0.5ml Eppendorf tube with holes poked by 18G needle. The 0.5ml Eppendorf Tube harboring the bones was then put into a 1.5ml Eppendorf tube with 100ml culture media. The bone marrow was isolated by centrifuge at 10,000g for 15s at 4Cº, followed by filtering the cells through 70μm mesh (Falcon) and lysis of red blood cells using ammonium-chloride-potassium (ACK) buffer (deionized water with 0.15M NH4Cl, 1mM KHCO3, and 0.1mM Na2EDTA). These cells were then i.v. injected into the irradiated (1100Rads) WT mice at the quantity of ∼4 million cells/mice. The chimera mice were sacrificed 10 weeks post reconstitution.

#### Broncho-alveolar lavage (BAL) fluid

After euthanization of the mouse with an overdose of Ketamine/xylazine, the trachea was exposed with incision on the skin and blunt dissection of the thyroid as well as muscle. An small incision was made between thyroid cartilage and Cricoid cartilage, followed by insertion of pipette tip (20ml tip topping on 1000ml tip). Using p1000 pipette, BAL fluid was obtained by flushing the airway five times with a single inoculum of 1000μl sterile PBS (with 2% FBS and 1X Pen/Strep/Glutamate). Cells in BAL fluid was spun down at 1600rpm for 5min followed by lysis of red blood cells using ammonium-chloride-potassium (ACK) buffer. The cells were then palleted (1600rpm for 5min) and resuspended for flow cytometry or cell culture.

#### Single cell suspension from Lung homogenate

Animals were euthanized with an overdose of Ketamine/xylazine. Lungs were perfused with 10ml PBS from the right ventricle of heart through pulmonary circulation. Following the harvest, lung tissue was minced and put in digestion buffer (IMDM with 183 U/ml type 2 collagenase (Worthington)). After 30min incubation at 37 Cº, the homogenate was put through the m_lung_02 program on a gentleMACS tissue disrupter (Miltenyi). (Specifically, for Spider-βGal staining, to avoid stress from the process, manual homogenization with metal mesh was used in this step). A 70μm mesh (Falcon) was used to filter the suspension. The flow through will be washed using MACS buffer (PBS with 1% FBS). The cells were then pelleted at 1600rpm for 5min, followed by lysis of red blood cells using ammonium-chloride-potassium (ACK) buffer (deionized water with 0.15M NH_4_Cl, 1mM KHCO_3_, and 0.1mM Na_2_EDTA). The single cell suspension would be utilized to perform flow cytometry or FACS sorting.

#### In vitro GM-CSF stimulation experiment

Cells were palleted from BAL washes and put in 12-well plate in complete medium (RPMI 1640 with 10% FBS and 1X Pen/Strep/Glutamate) at 37 Cº in 5% CO_2_. The non-adherent cells were washed off with warm PBS after 2 hours, and AMs would be purified from these cells by its adherent nature. The remaining adherent cells were cultured overnight in complete medium without GM-CSF, followed by culture in complete media supplemented with 10ng/ml (unless specified) GM-CSF for 24 hours. AMs were then detached with 0.25% Tyrosine-EDTA solution (Gibco) and put through flow cytometry to evaluate its proliferation.

#### In vitro inhibition of CBFβ-RUNX1 interaction

Cells of BAL fluid and bone marrow were harvested from the same mice. AMs was purified in 12-well plates as described in the last section. Followed overnight culture in complete medium without GM-CSF, AMs were treated with 100μM, 50μM or 0μM Ro 5-3335 (TargetMol) in complete medium supplemented with 10ng/ml GM-CSF for 24 hours.

Bone Marrow was isolated as previously described^59^. After filtering the cells through 70μm mesh (Falcon), lysis of red blood cells was performed with ACK buffer. The cells were then resuspended in culture media (DMEM with 50ng/ml M-CSF, and 1X Pen/Strep/Glutamate) and seed in the 10cm Petri dish (Falcon). Counting the date of seeding as day0, cells was culture at 37 Cº with 5% CO_2_ for 7 days with change for fresh media on day4. The BMDMs were detached and re-seed in 12-well plates on day7. After resting the cells in culture media overnight, BMDMs were treated with 100μM, 50μM or 0 μM Ro 5-3335 in culture media for 24 hours.

0.25% Trypsin-EDTA solution (Gibco) was then used to detach AMs/BMDMs from the plate, followed by evaluation of proliferation using flow cytometry.

#### Colony formation assay

Colony formation assays were performed with cells in BAL from indicated groups as previously described^5^. Briefly, freshly isolated cells from BAL were seeded at 20,000 cells in 35 mm culture dishes. These cells were then cultured in MethoCult medium (M3231, Stem Cell Technologies) supplemented with 10 ng/ml GM-CSF, 50 mg/ml penicillin/streptomycin and 2 mM glutamine. The number of colonies were counted on day 19 and day 21 after plating.

#### Flow Cytometry

Detached AMs or cells from lung homogenate were prepared for flow cytometry. The information of Antibodies used can be found In **Supplementary Table1**. The Fc receptor was blocked with α-mouse CD16/CD32 (BioXcell). Cells were then incubated in solutions with antibodies targeting surface makers on ice for 30min. If no intracellular protein of interest is involved, the samples would be ready after washing off the antibodies.

If intracellular staining is needed, after staining for surface markers, cells were washed with MACS buffer twice. The cells be fixed with fixation butter (eBioscience) at room temperature for 30min, followed by 1h incubation in permeabilization buffer (eBioscience) on ice after washing off the fixation buffer. And staining of intracellular markers were then performed on ice for 1h. The samples would be ready after washing off the antibodies.

Samples were put in MACS buffer and put through FACS Attune. Analysis of the data was enabled with FlowJo.

#### Brdu incorporation

BrdU (Sigma, 1mg/mouse in sterile PBS) was i.p. injected 1 day before harvest. And flow cytometry was used to evaluate the Ki-67 and BrdU incorporation of AMs.

#### SPiDER-bGal Staining

The cells in the single cell suspension from lung homogenate were stained with extracellular markers for flow before putting through SPiDER-bGal (Dojindo) staining. Cells were palleted and resuspend in 1ml warn RPMI-1640 with Bafilomycin A1 (1:1000), which was then incubated at 37 Cº in 5% CO_2_ for 1h. Another 1ml warm RPMI-1640 with SPiDER-bGal (1:1000, the final concentration in the working solution is 1:2000) and Bafilomycin A1 (1:1000) was added in. And the stain would be wash off after another 30min incubation 37 Cº in 5% CO_2_. These cells were stained with Zombie dye (Biolegend) to evaluate the viability of the cell. For quantification of SPiDER-bGal staining, geometry mean was taken in the live AMs (Zombie-CD11c+Siglec F+).

#### Nuclear Extraction and Western Blot

NE-PER™ Nuclear and Cytoplasmic Extraction Reagents (ThermoFisher) was used for nuclear extraction of AMs. Following the protocol, around 0.5*10^6^ Cells from BAL was spin down and resuspend in 50ml pre-cooled CRE I solution by vertexing for 15s. After 10min incubation on ice, 2.75ml pre-cooled CRE II solution was added in. Start and end with 5s vertexing, 1 min incubation was done on ice, followed by 5 minutes at maximum speed in a microcentrifuge at 4Cº. The supernatant (cytoplasmic extract) was transferred to pre-chilled Eppendorf Tube on ice. 25ml pre-cooled NER solution was used to resuspend the pallet by repeated vertexing for 15s every 10min for a total 40min. 10min centrifuge at the highest speed (4Cº) was then performed, and the supernatant (nuclear extract) was collected in pre-chilled Eppendorf Tube.

Denature of Protein of both the cytoplasmic extract and nuclear extract was performed utilizing 4X Protein Sample Loading Buffer (LI-COR) with β-mercaptoethanol. Samples were boiled at 100Cº for 5min. Electrophoresis was performed with pre-casted NuPAGE® Novex 4-12% Bis-Tris gel (Invitrogen) in MES buffer. Transfer was done overnight in transfer buffer (1XTris/Glycine buffer with 10% methanol) at 30V, 4Cº. After the transfer, total protein was stained using REVERT™ Total Protein Stain (LI-COR). After imaging the membrane in the 700nm channel in Odyssey^®^ imaging system, protein stain was washed off using destaining solution in the kit. The membrane was put in Intercept® (TBS) Blocking Buffer (LI-COR) for 24h to ensure elimination of effect for the destaining solution. Blocking was done using the same buffer, and the membrane was incubated in primary antibody of CBFβ (Abcam, 1:1000) overnight. Secondary antibody incubation was performed in room temperature in Blocking buffer with anti-Rabbit antibody (LI-COR, 1:15000) 0.1% Tween 20 and 0.01%SDS. After washing the membrane in PBST (PBS +0.1% Tween20), the membrane was dried in dark at room temperature overnigh. And the membrane was then visualized in 800nm channel in Odyssey^®^ imaging system. ImageStudio was utilized for quantification. Expression of CBFβ was calculated as:

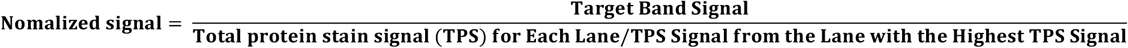

#### Quantitative RT-PCR

RNA extraction of freshly isolated cells in BAL or detached AMs was performed using total RNA purification kit (Sigma) together with DNase I (invitrogen) following the protocol in the kit. Reverse transcription for cDNA was then executed with Random primers (Invitrogen) and Moloney murine leukemia virus (MMLV) reverse transcriptase (Invitrogen), followed by RT-PCR with Fast SYBR Green PCR Master Mix (Applied Biosystems) in QuantStudio3 (Applied Bioscience). Relative expression of the target gene was calculated with respect to house-keeping gene, hypoxanthine phosphoribosyl transferase (Hprt).

#### scRNAseq analysis

The primary data was annotated and aligned using CellRanger 4.0. Using R project and the package of Seurat, we were able to further process the data. Genes that have been detected in at least 3 cells were included. And cells that has 200-4000 total genes detected, with less than 5% mitochondria genes were considered to pass quality control. The clustering was then performed as previously described^30^ and the vignette can be found in https://satijalab.org/seurat/articles/pbmc3k_tutorial.html. Of note, since the number AMs from the young mice were almost twice as many as that of AMs from the aged mice, the same number of AMs from the young mice was randomized out for the combined analysis to ensure the equal input in finding the differential expressed genes.

Contaminating myeloid cells were excluded as described in the manuscript. And clustering was performed again using the AM population. Module score was calculated in using the AddModuleScore() function of Seurat with gene lists indicated. Trajectory analysis was performed using Monocle3 in conjunction with Signac, an extension of Seurat, as described in the paper^31,60^, and the vignette can be found in https://satijalab.org/signac/articles/monocle.html. The proliferating cells were chosen as the “stem cells”. For pathway analysis, 1000 cells from each group were randomized out and put through Gene Set Enrichment Analysis as described^37,38^.

For human data, publicly available dataset (GSE 128033) was utilized. AM population was defined as described in the manuscript. The integration of Young, Middle-Aged and Aged group was then performed using the graph-based method described^61^ to strike a balance between the resolution of cell population and the limited AM number in the Aged group.

## Notes

### Competing Interest Statement

The authors have declared no competing interest.

## Reference

1. Mowat, A. M. I., Scott, C. L. & Bain, C. C. Barrier-tissue macrophages: Functional adaptation to environmental challenges. Nat. Med. 23, 1258–1270 (2017).

2. Hussell, T. & Bell, T. J. Alveolar macrophages: Plasticity in a tissue-specific context. Nat. Rev. Immunol. 14, 81–93 (2014).

3. Guilliams, M. et al. Alveolar macrophages develop from fetal monocytes that differentiate into long-lived cells in the first week of life via GM-CSF. J. Exp. Med. 210, 1977–92 (2013).

4. Evren, E. et al. CD116+ fetal precursors migrate to the perinatal lung and give rise to human alveolar macrophages. J. Exp. Med. 219, (2022).

5. Soucie, E. L. et al. Lineage-specific enhancers activate self-renewal genes in macrophages and embryonic stem cells. Science (80-.). 351, (2016).

6. Hashimoto, D. et al. Tissue-resident macrophages self-maintain locally throughout adult life with minimal contribution from circulating monocytes. Immunity 38, 792–804 (2013).

7. Nayak, D. K. et al. Long-Term Persistence of Donor Alveolar Macrophages in Human Lung Transplant Recipients That Influences Donor-Specific Immune Responses. Am. J. Transplant. 16, 2300–2311 (2016).

8. Draijer, C., Penke, L. R. K. & Peters-Golden, M. Distinctive Effects of GM-CSF and M-CSF on Proliferation and Polarization of Two Major Pulmonary Macrophage Populations. J. Immunol. 202, 2700–2709 (2019).

9. Yu, X. et al. The Cytokine TGF-β Promotes the Development and Homeostasis of Alveolar Macrophages. Immunity 47, 903-912.e4 (2017).

10. Schneider, C. et al. Induction of the nuclear receptor PPAR-γ by the cytokine GM-CSF is critical for the differentiation of fetal monocytes into alveolar macrophages. Nat. Immunol. 15, 1026–1037 (2014).

11. Rauschmeier, R. et al. Bhlhe40 and Bhlhe41 transcription factors regulate alveolar macrophage self-renewal and identity. EMBO J. 1–20 (2019). doi:10.15252/embj.2018101233

12. Nakamura, A. et al. Transcription repressor Bach2 is required for pulmonary surfactant homeostasis and alveolar macrophage function. 210, 2191–2204 (2013).

13. Suwankitwat, N. et al. The actin-regulatory protein Hem-1 is essential for alveolar macrophage development. 218, (2021).

14. Mccowan, J. et al. The transcription factor EGR2 is indispensable for tissue-specific imprinting of alveolar macrophages in health and tissue repair. 2, 1–44 (2021).

15. Zhu, B. et al. Uncoupling of macrophage inflammation from self-renewal modulates host recovery from respiratory viral infection. Immunity 54, 1200-1218.e9 (2021).

16. Gao, X. et al. TFAM-Dependent Mitochondrial Metabolism Is Required for Alveolar Macrophage Maintenance and Homeostasis. J. Immunol. 208, 1456–1466 (2022).

17. Herranz, N., Gil, J., Herranz, N. & Gil, J. Mechanisms and functions of cellular senescence. JCI J. Clin. Investig. 128, 1238–1246 (2018).

18. Hernandez-Segura, A., Nehme, J. & Demaria, M. Hallmarks of Cellular Senescence. Trends Cell Biol. 28, 436–453 (2018).

19. Yousefzadeh, M. J. et al. An aged immune system drives senescence and ageing of solid organs. Nature 594, (Springer US, 2021).

20. Moliva, J. I. et al. Molecular composition of the alveolar lining fluid in the aging lung. Age (Omaha). 36, 1187–1199 (2014).

21. Wroblewska, L. et al. Aging impairs alveolar macrophage phagocytosis and increases influenza-induced mortality in mice. J. Immunol. 199, 1060–1068 (2017).

22. McQuattie-Pimentel, A. C. et al. The lung microenvironment shapes a dysfunctional response of alveolar macrophages in aging. J. Clin. Invest. 131, (2021).

23. Speck, N. A. & Gilliland, D. G. Core-binding factors in haematopoiesis and leukaemia. Nat. Rev. Cancer 2, 502–513 (2002).

24. Wang, Q. et al. The CBFbeta subunit is essential for CBFalpha2 (AML1) function in vivo. Cell 87, 697–708 (1996).

25. Kundu, M. et al. Role of Cbfb in hematopoiesis and perturbations resulting from expression of the leukemogenic fusion gene Cbfb-MYH11. Blood 100, 2449–2456 (2002).

26. Kundu, M. & Liu, P. P. CBFβ is involved in maturation of all lineages of hematopoietic cells during embryogenesis except erythroid. 30, 164–169 (2003).

27. Murakami, K. et al. A RUNX–CBFβ-driven enhancer directs the Irf8 dose-dependent lineage choice between DCs and monocytes. Nat. Immunol. 22, (2021).

28. Cardani, A., Boulton, A., Kim, T. S. & Braciale, T. J. Alveolar Macrophages Prevent Lethal Influenza Pneumonia By Inhibiting Infection Of Type-1 Alveolar Epithelial Cells. PLoS Pathog. 13, 1–25 (2017).

29. Mould, K. J. et al. Airspace macrophages and monocytes exist in transcriptionally distinct subsets in healthy adults. Am. J. Respir. Crit. Care Med. 203, 946–956 (2021).

30. Satija, R., Farrell, J. A., Gennert, D., Schier, A. F. & Regev, A. Spatial reconstruction of single-cell gene expression data. Nat. Biotechnol. 33, 495–502 (2015).

31. Trapnell, C. et al. The dynamics and regulators of cell fate decisions are revealed by pseudotemporal ordering of single cells. Nat. Biotechnol. 32, 381–386 (2014).

32. Wong, D. J. et al. Module Map of Stem Cell Genes Guides Creation of Epithelial Cancer Stem Cells. Cell Stem Cell 2, 333– 344 (2008).

33. Kowalczyk, M. S. et al. Single-cell RNA-seq reveals changes in cell cycle and differentiation programs upon aging of hematopoietic stem cells. Genome Res. 25, 1860–1872 (2015).

34. Heng, T. S. P., Painter, M. W., Immunological, T. & Project, G. The Immunological Genome Project : networks of gene expression in immune cells. 9, 1091–1094 (2008).

35. Zhu, B., Wu, Y. & Huang, S. Uncoupling of macrophage inflammation from self-renewal modulates host recovery from respiratory viral infection Uncoupling of macrophage inflammation from self-renewal modulates host recovery from respiratory viral infection. Immunity 54, 1–19 (2021).

36. Shi, J., Hua, L., Harmer, D., Li, P. & Ren, G. Cre Driver Mice Targeting Macrophages. Methods Mol Biol. 1784, 263–275 (2018).

37. Subramanian, A. et al. Gene set enrichment analysis: A knowledge-based approach for interpreting genome-wide expression profiles. Proc. Natl. Acad. Sci. U. S. A. 102, 15545–15550 (2005).

38. Daly, M. J. et al. PGC-1α-responsive genes involved in oxidative phosphorylation are coordinately downregulated in human diabetes. Nat. Genet. 34, 267–273 (2003).

39. Chatsirisupachai, K., Palmer, D., Ferreira, S. & de Magalhães, J. P. A human tissue-specific transcriptomic analysis reveals a complex relationship between aging, cancer, and cellular senescence. Aging Cell 18, 1–5 (2019).

40. Hattori, N. et al. Bleomycin-induced pulmonary fibrosis in fibrinogen-null mice Find the latest version : in fibrinogen-null mice. J. Clin. Invest. 106, 1341–1350 (2000).

41. García-Prieto, E. et al. Resistance to bleomycin-induced lung fibrosis in MMP-8 deficient mice is mediated by interleukin-10. PLoS One 5, (2010).

42. Jiang, C. et al. Serpine 1 induces alveolar type II cell senescence through activating p53-p21-Rb pathway in fibrotic lung disease. Aging Cell 16, 1114–1124 (2017).

43. Khan, S. S. et al. A null mutation in SERPINE1 protects against biological aging in humans. Sci. Adv. 3, (2017).

44. Samarakoon, R., Higgins, S. P., Higgins, C. E. & Higgins, P. J. The TGF-β1/p53/PAI-1 signaling axis in vascular senescence: Role of Caveolin-1. Biomolecules 9, 1–12 (2019).

45. Lee, B. Y. et al. Senescence-associated β-galactosidase is lysosomal β-galactosidase. Aging Cell 5, 187–195 (2006).

46. Adams, P. D. et al. Lysosome-mediated processing of chromatin in senescence. J. Cell Biol. 202, 129–143 (2013).

47. Schafer, M. J. et al. Cellular senescence mediates fibrotic pulmonary disease. Nat. Commun. 8, (2017).

48. Calhoun, C. et al. Senescent Cells Contribute to the Physiological Remodeling of Aged Lungs. Journals Gerontol. -Ser. A Biol. Sci. Med. Sci. 71, 153–160 (2016).

49. Lee, S. et al. Molecular programs of fibrotic change in aging human lung. Nat. Commun. 12, 1–10 (2021).

50. Lee, J. et al. Molecular programs of fibrotic change in aging human lung. (2021).

51. Morse, C. et al. Proliferating SPP1/MERTK-expressing macrophages in idiopathic pulmonary fibrosis. Eur. Respir. J. 54, (2019).

52. Gschwend, J. et al. Alveolar macrophages rely on GM-CSF from alveolar epithelial type 2 cells before and after birth. J. Exp. Med. 218, (2021).

53. Westphalen, K. et al. Sessile alveolar macrophages communicate with alveolar epithelium to modulate immunity. Nature 506, 503–506 (2014).

54. Svedberg, F. R. et al. The lung environment controls alveolar macrophage metabolism and responsiveness in type 2 inflammation. Nat. Immunol. 20, 571–580 (2019).

55. Subramanian, S. et al. Long-term culture-expanded alveolar macrophages restore their full epigenetic identity after transfer in vivo. Nat. Immunol. (2022). doi:10.1038/s41590-022-01146-w

56. Liu, Z. et al. Fate Mapping via Ms4a3-Expression History Traces Monocyte-Derived Cells. Cell 178, 1509-1525.e19 (2019).

57. Omori, S. et al. Generation of a p16 Reporter Mouse and Its Use to Characterize and Target p16high Cells In Vivo. Cell Metab. 1–15 (2020). doi:10.1016/j.cmet.2020.09.006

58. Amend, S. R., Valkenburg, K. C. & Pienta, K. J. Murine Hind Limb Long Bone Dissection and Bone Marrow Isolation. J. Vis. Exp. 110, 3–6 (2016).

59. Aurrand-Lions, M. & Mancini, S. J. C. Murine bone marrow niches from hematopoietic stem cells to B cells. Int. J. Mol. Sci. 19, 1–18 (2018).

60. Qiu, X. et al. Reversed graph embedding resolves complex single-cell trajectories. Nat. Methods 14, 979–982 (2017).

61. Stuart, T. et al. Comprehensive Integration of Single-Cell Data. Cell 177, 1888-1902.e21 (2019).

