## Supplemental Figures for "Single cell RNA sequencing reveals mechanisms underlying a senescence-like phenotype of Alveolar Macrophages during Aging"

Supplementary Figure 1

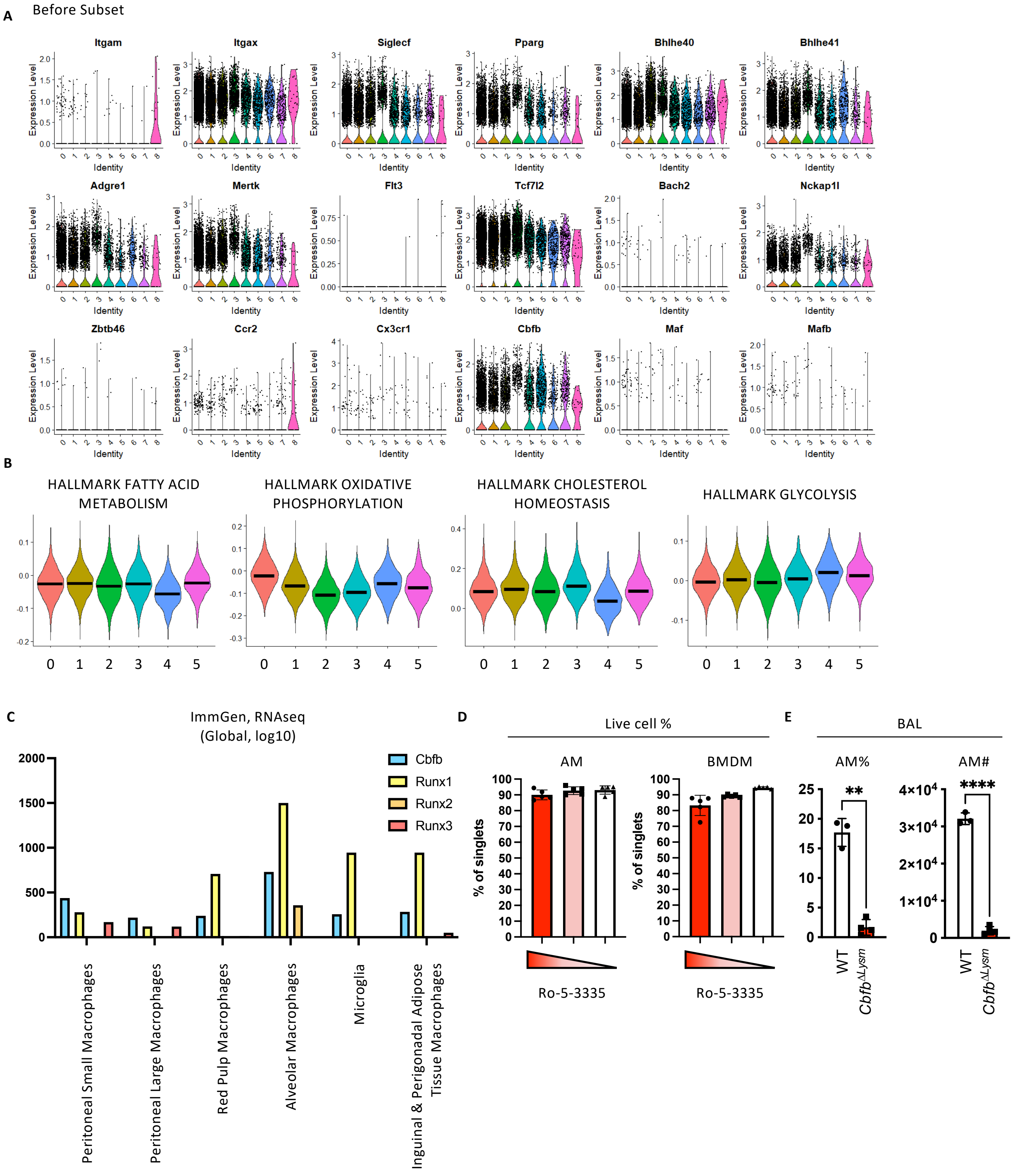

Supplementary Figure 2

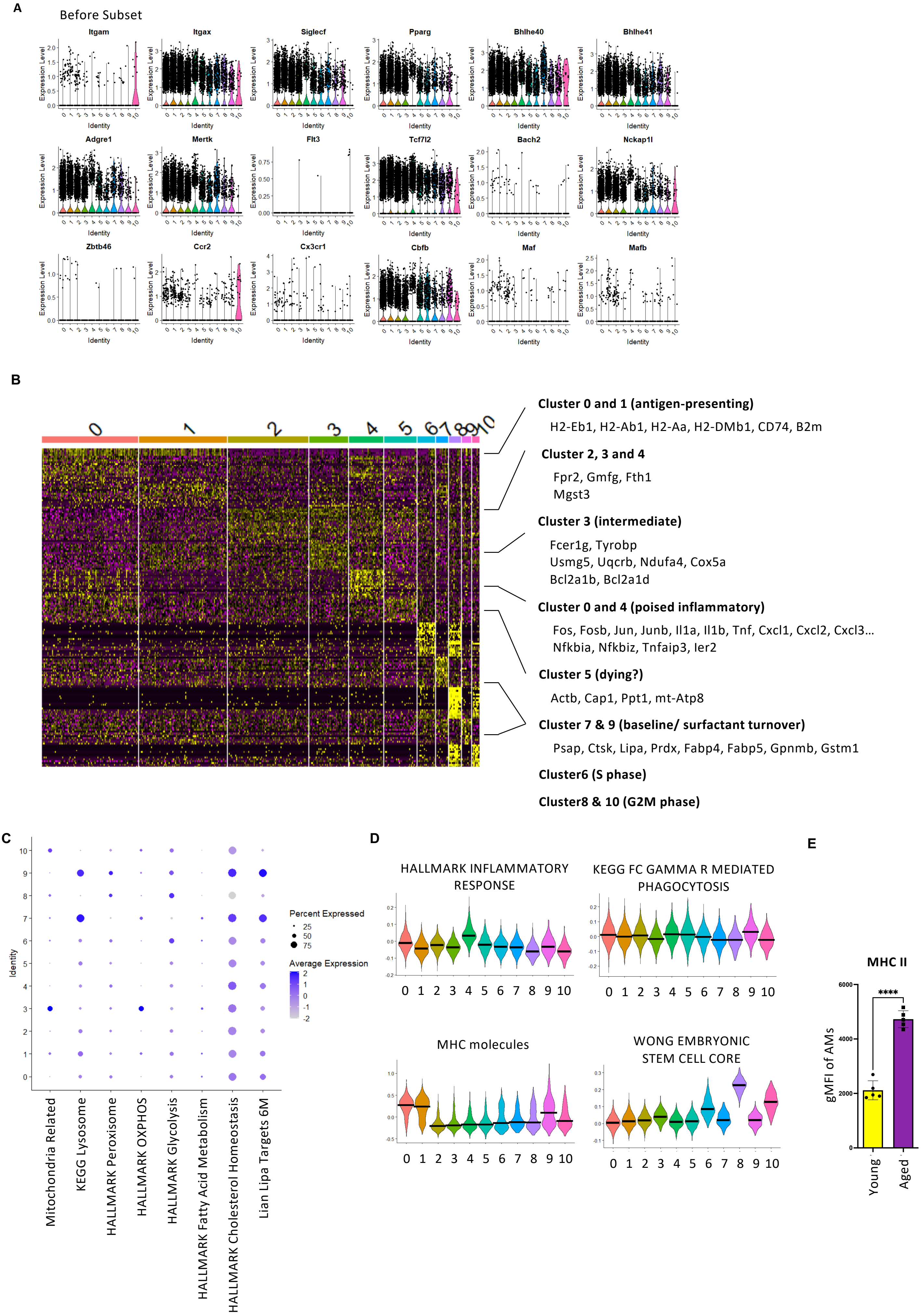

Supplementary Figure 3

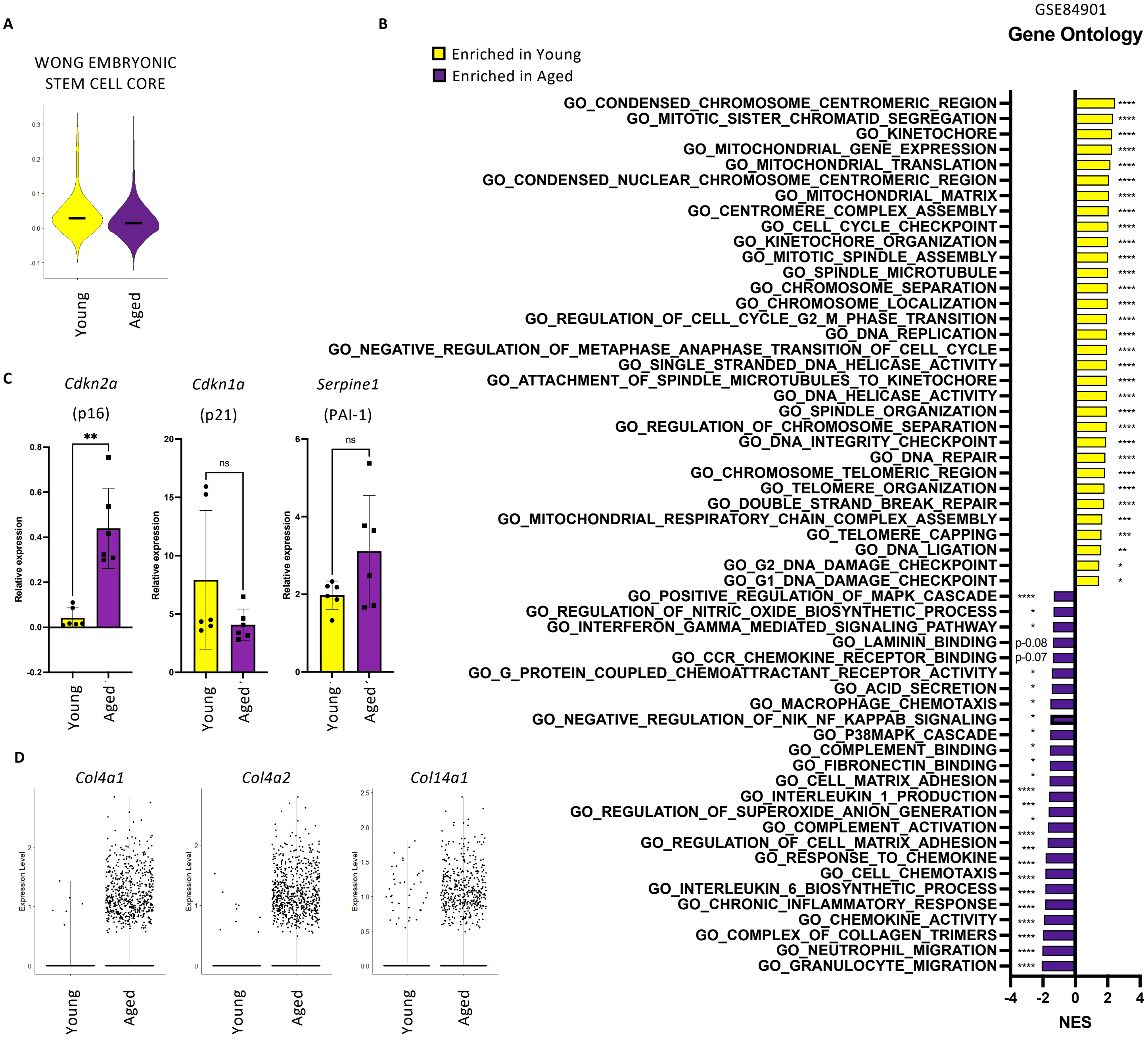

Supplementary Figure 4

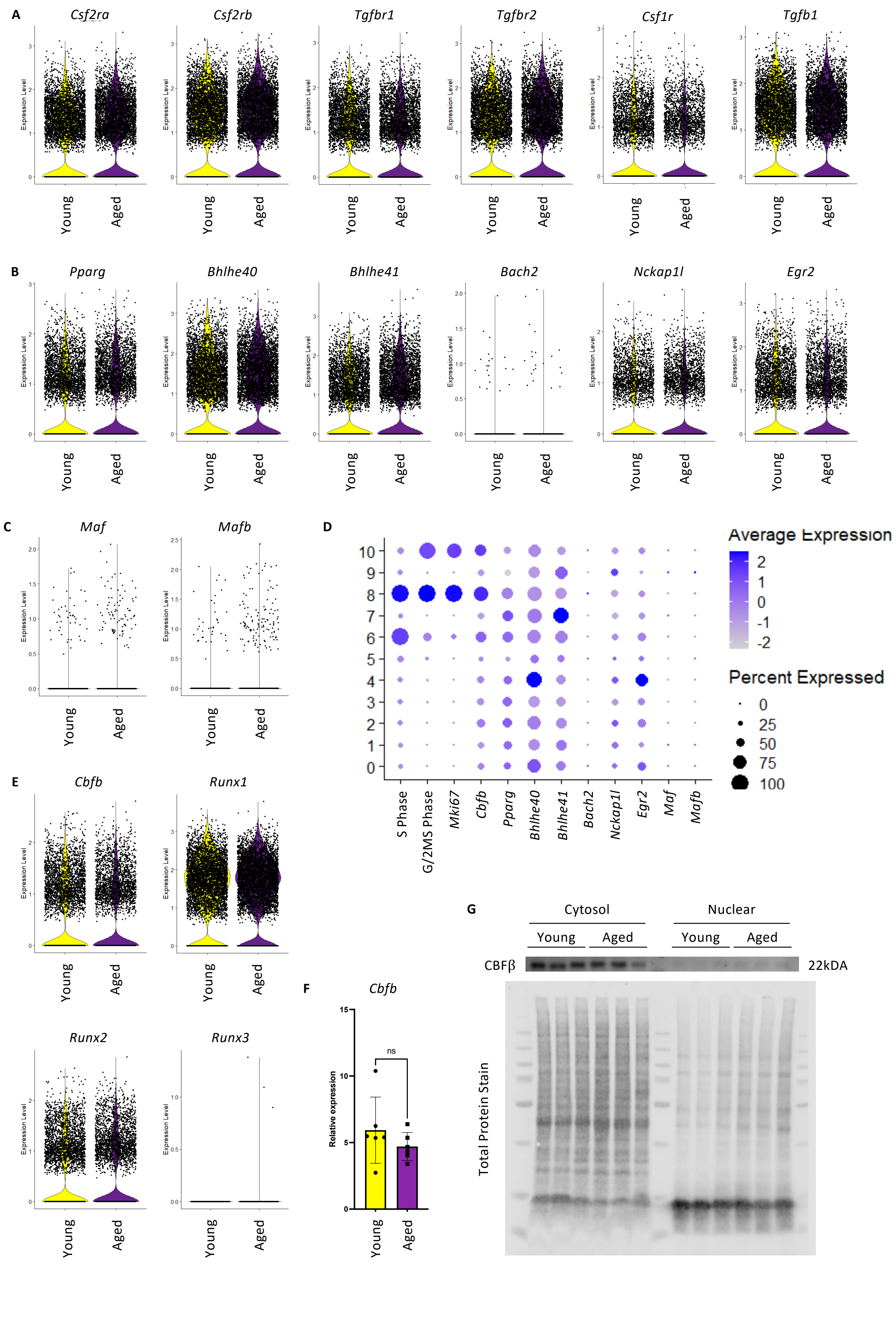

Supplementary Figure 5

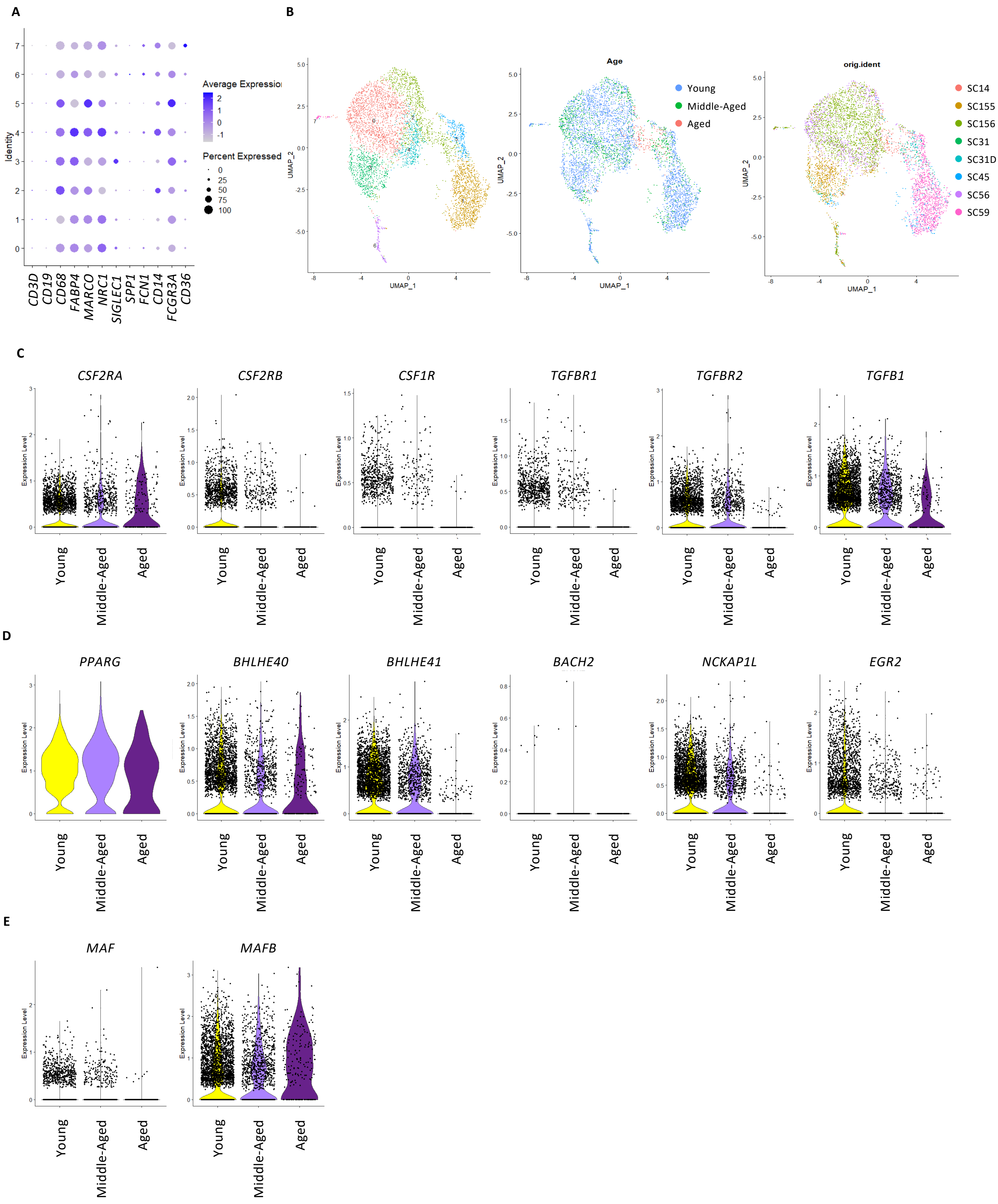
